# Nanaerobic growth enables direct visualization of dynamic cellular processes in human gut symbionts

**DOI:** 10.1101/2020.05.22.111492

**Authors:** Leonor García-Bayona, Michael J. Coyne, Noam Hantman, Paula Montero-Llopis, Salena Von, Takeshi Ito, Michael H. Malamy, Marek Basler, Blanca Barquera, Laurie E. Comstock

## Abstract

Mechanistic studies of anaerobic gut bacteria have been hindered by the lack of a fluorescent protein system to track and visualize proteins and dynamic cellular processes in actively growing bacteria. Although underappreciated, many gut “anaerobes” are able to respire using oxygen as the terminal electron acceptor. The oxygen continually released from gut epithelial cells creates an oxygen gradient from the mucus layer to the anaerobic lumen (1), with oxygen available to bacteria growing at the mucus layer. Using a combination of analyses, we show that *Bacteroides* species are metabolically and energetically robust and do not mount stress responses in the presence of 0.10 - 0.14% oxygen, defined as nanaerobic conditions (2). Taking advantage of this metabolic capability, we show that nanaerobic growth provides sufficient oxygen for the maturation of oxygen-requiring fluorescent proteins in *Bacteroides* species. Type strains of four different *Bacteroides* species show bright GFP fluorescence when grown nanaerobically versus anaerobically. We compared four different red fluorescent proteins and found that mKate2 yields high fluorescence intensity in our assay. We show that GFP-tagged proteins can be localized in nanaerobically growing bacteria. In addition, we used time-lapse fluorescence microscopy to image dynamic Type VI secretion system processes in metabolically active *B. fragilis*. The ability to visualize fluorescently-labeled *Bacteroides* and fluorescently-linked proteins in actively growing nanaerobic gut symbionts ushers in a new age of imaging analyses in these bacteria.

**Significance:** Despite many recent technological advances to study the human gut microbiota, we still lack a facile system to image dynamic cellular processes in most abundant gut species due to the requirement of oxygen for chromophore maturation of commonly used fluorescent proteins. Here, we took advantage of the ability of anaerobes of the gut microbiota to respire aerobically and grow robustly at 0.10– 0.14% oxygen. This physiologic concentration of oxygen is sufficient for fluorescent proteins to mature, allowing for visualization of biological processes never before imaged in these bacteria. This advance will allow for numerous types of analyses in actively-growing “nanaerobic” gut bacteria including subcellular protein localizations, single-cell analyses, biofilm imaging, and protein interactions with other microbes and the host.

## INTRODUCTION

The rapid increase in knowledge of the human gut microbiota and its impact on health and disease processes is the result of a multi-disciplinary approach from microbiology, immunology, ecology, genomics, computational biology and metabolomics, among others. The genetic toolkit to analyze bacterial members of this ecosystem has also greatly expanded in recent years (3-8) allowing better and faster analyses and mechanistic studies. Despite these advances, we have still been unable to use GFP and other oxygen requiring fluorescent proteins to visually track proteins or to visualize dynamic cellular processes in most gut symbionts.

The optimization of the flavin mononucleotide (FMN)-based fluorescent protein for use in anaerobic organisms brought hope for the development of a GFP equivalent for anaerobic microbes (9). Indeed, FMN-based fluorescent proteins have been successfully applied to studies of some anaerobes (10-12), but few studies have reported successful use of these or other fluorescent proteins in most anaerobic gut bacteria, especially in actively growing bacteria. Another approach to fluorescently label gut anaerobes used click chemistry to label surface polysaccharides (13). Bacteria metabolically labeled *in vitro* were tracked *in vivo* for up to 12 hours before the fluorescent signal was lost from the surface. This labeling allowed imaging of polysaccharide association with immune cells (13).

Other fluorescent imaging techniques have been applied to study the spatial structure of bacterial communities. Advances in fluorescent *in situ* hybridization (FISH) are illuminating the architecture of synthetic and natural microbial communities (14, 15), with the recent application of multiplexed fluorescence spectral imaging to image the spatial ecology of numerous different taxa of bacteria of the human tongue (16). A fluorescent protein-based analysis also revealed community structure in the mammalian gut (5). In this study, six different *Bacteroides* species were differentiated *in vivo* using constructs that resulted in three different levels of expression of GFP and mCherry by quantification of single-cell fluorescent profiles.

However, we still lack a fluorescent protein-based system to study dynamic processes in most actively growing gut anaerobes (reviewed (17)). The human intestinal microbiota is dominated by bacteria that are considered to be strict anaerobes. Almost half of the gut bacteria in most healthy humans are members of the Gram negative order Bacteroidales. These include numerous species of *Bacteroides, Parabacteroides, Prevotella, Alistipes*, and others. The gut microbiota also contains a large proportion of members of the phylum Firmicutes, which includes many diverse families. Some, such as the members of the *Clostridiaceae* family, are strict anaerobes, while others, such as members of the Lactobacillaceae family, have varying abilities to grow in room air. In addition, there are abundant genera of other phyla including Actinobacteria and Verrucomicrobia with varying abilities to grow in the presence of oxygen.

Cytochrome *bd* is one of the terminal oxidases of aerobic respiration in prokaryotes (reviewed (18)). In contrast to many heme-copper oxidases that are typically expressed when bacteria are grown in high aeration, cytochrome *bd* is usually expressed in low oxygen conditions. This likely reflects the fact that cytochrome *bd* has a much higher affinity for oxygen than most heme-copper oxidases.(19, 20). Cytochrome *bd* accepts electrons from quinols and donates them to oxygen to make water. The enzyme generates an electrochemical proton gradient by a membrane charge separation. In this way, cytochrome *bd* is able to conserve the energy from nanaerobic respiration for use by the cell (21).

Among the abundant gut bacteria, *Bacteroides* species and *Akkermansia mucinophila* have been shown to have cytochrome *bd* and to respire using oxygen as the terminal electron acceptor (2, 22). This metabolic capability was shown in both *B. fragilis* and *A. mucinophila* to lead to the consumption of oxygen during nanaerobic growth. In this study, we show that the nanaerobic oxygen concentration under which *Bacteroides* species likely grow and thrive when at the mucus layer is sufficient to allow for the maturation of commonly used fluorescent proteins in order to image active processes in healthy, metabolically active bacterial cells.

## RESULTS

### Analysis of *B. fragilis* grown under nanaerobic conditions

Four commonly studied *Bacteroides* strains, *B thetaiotaomicron* VPI-5482, *B. ovatus* ATCC 8483, *B. vulgatus* ATCC 8482 and *B. fragilis* 638R, all grow robustly in a nanaerobic atmosphere (0.1% - 0.14% or 1000 – 1400 ppm atmospheric oxygen) as previously shown (2), reaching a maximal OD_600_ similar to that reached during anaerobic growth (Fig 1A, Fig S1). During anaerobic respiration in *Bacteroides*, fumarate reductase is the terminal component of the pathway, transferring electrons from quinols to fumarate. To show a benefit from nanaerobic growth, we made a *B. fragilis* mutant deleted for the three fumarate reductase genes *frdABC*. The Δ*frd* mutant is severely defective for growth under anaerobic conditions, but its growth is improved under nanaerobic conditions (Fig 1A, (2)).

**Figure 1.**
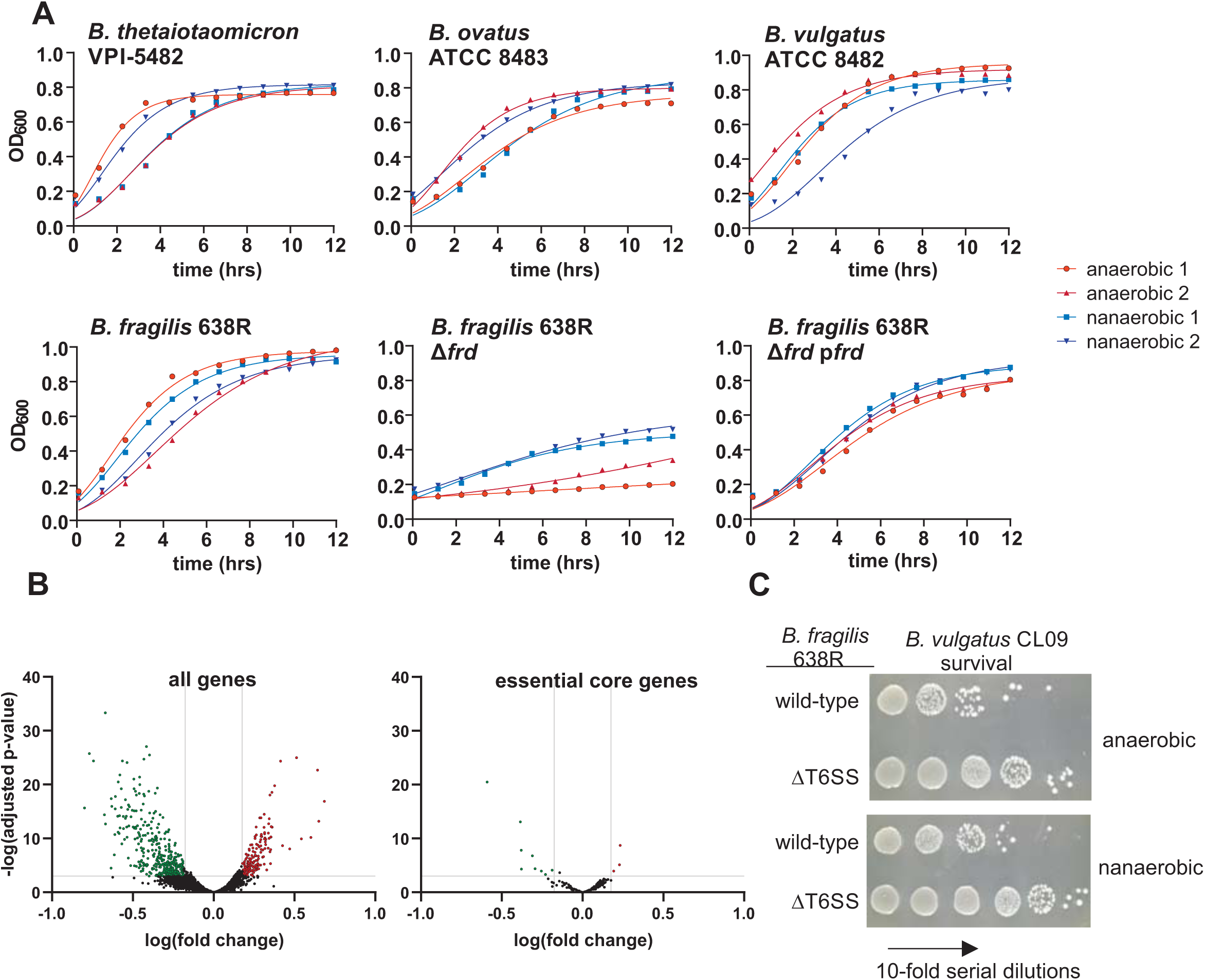
Analysis of nanaerobic growth properties of *Bacteroides*. **A**. Growth curves of four *Bacteroides* species and the *B. fragilis* Δ*frd* mutant and complemented mutant Δ*frd* p*frd* under anaerobic and nanaerobic conditions. **B**. Volcano plot showing differential expression of the 4332 genes of *B. fragilis* 638R (left panel). Volcano plot showing relative expression of the 372 core essential genes of *B. fragilis* under nanaerobic compared to anaerobic growth (right panel). Green dots represent genes significantly downregulated under nanaerobic conditions and red genes represent upregulated genes (significance defined as at least 1.5 fold and adjusted *p*-value of less than or equal to 0.001 by each of two measures, see text). **C**. Analysis of the ability of the *B. fragilis* GA3 T6SS to antagonize sensitive strain *B. vulgatus* CL09T03C10 under both anaerobic and nanaerobic conditions.

To determine potential differences in the physiology of *Bacteroides* when cultured with or without nanaerobic concentrations of oxygen, we performed RNASeq analysis of *B. fragilis* 638R. Using a strict cutoff for differential expression of at least 1.5 fold and an adjusted *p*-value less than or equal to 0.001, of the total 4332 genes, there were 171 genes upregulated and 405 genes downregulated under nanaerobic growth compared to anaerobic growth (Fig 1B, Table S1). Most categories of genes based on Clusters of Orthologous Groups of proteins (COG) assignment were split between being more highly expressed under anaerobic or nanaerobic conditions (Fig S2). For example, 13 *susCD* genes involved in the transport of carbohydrates and other nutrients across the outer membrane are upregulated under nanaerobic conditions. Under anaerobic conditions, 17 distinct *susCD* genes are more highly expressed. In terms of transcriptional regulators, 18 genes of at least 6 different families are more highly expressed under anaerobic conditions and six are more highly expressed under nanaerobic conditions (Table S1).

We next analyzed the RNASeq data to determine if the bacteria mount an oxidative stress response when grown nanaerobically. A prior study identified the *B. fragilis* transcriptional responses when exposed to room oxygen, 5% oxygen, or H_2_O_2_ (23). In that study, 30 genes classified as involved in stress responses were highly upregulated under at least one of the three stress conditions. Of these 30 genes, only three were significantly upregulated during nanaerobic growth, and 13 were significantly downregulated (Table 1). Genes encoding catalase and superoxide dismutase, two enzymes that are involved reactive oxygen species (ROS) response, were not differentially expressed between the two conditions.

**Table 1.**
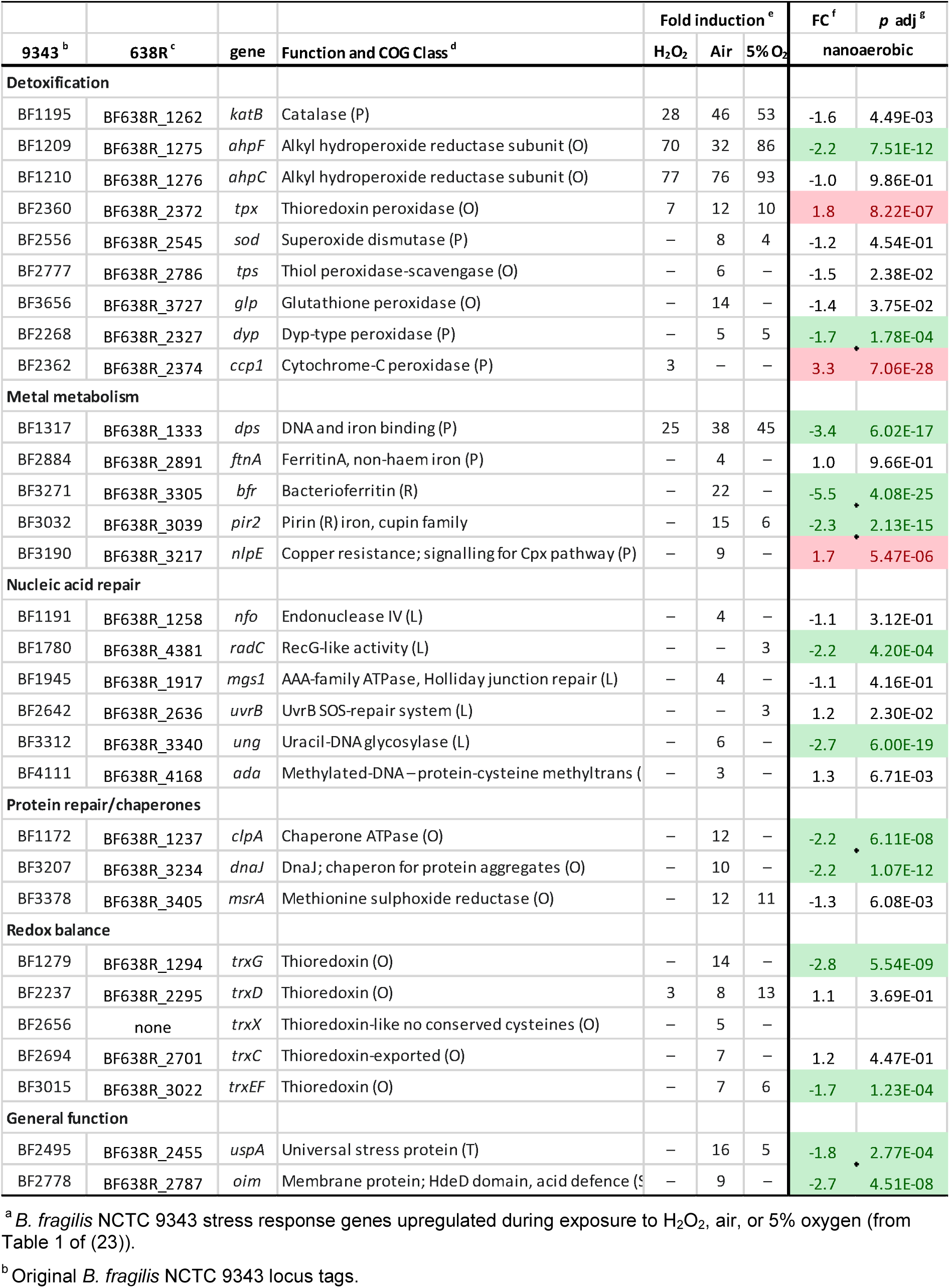
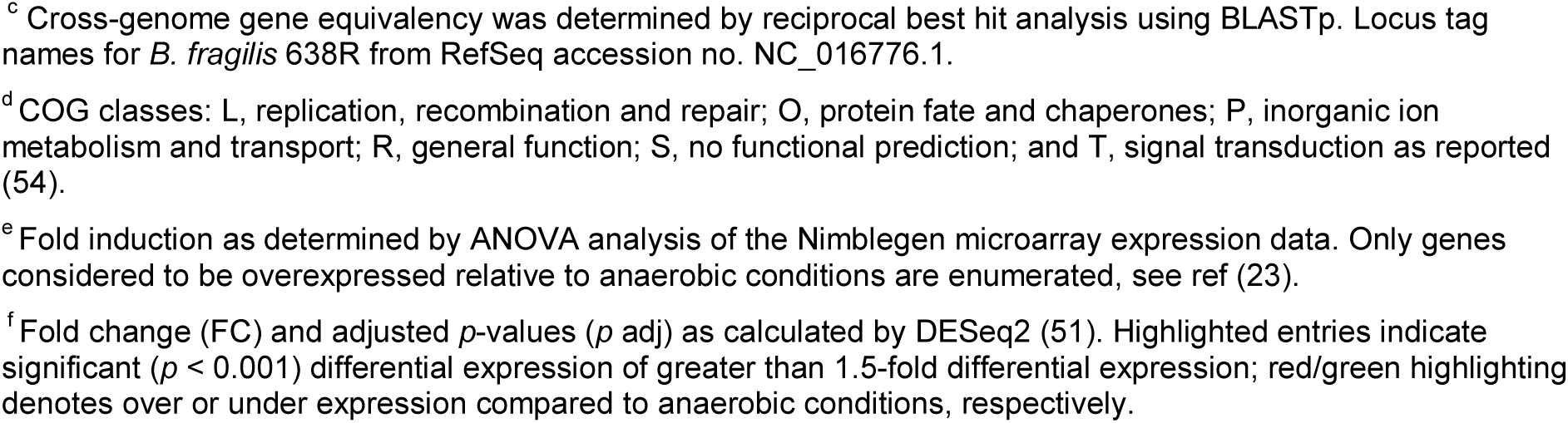
Comparison of differentially expressed genes under oxidative stress as previously reported (ref (23)) versus nanaerobic growth

|  |  |  |  | Fold induction<sup>e</sup> |  |  | FC<sup>f</sup> | p adj<sup>g</sup> |
| --- | --- | --- | --- | --- | --- | --- | --- | --- |
| 9343<sup>b</sup> | 638R<sup>c</sup> | gene | Function and COG Class<sup>d</sup> | H<sub>2</sub>O<sub>2</sub> | Air | 5% O<sub>2</sub> | nanoaerobic |  |
| Detoxification |  |  |  |  |  |  |  |  |
| BF1195 | BF638R_1262 | *katB* | Catalase (P) | 28 | 46 | 53 | -1.6 | 4.49E-03 |
| BF1209 | BF638R_1275 | *ahpF* | Alkyl hydroperoxide reductase subunit (O) | 70 | 32 | 86 | -2.2 | 7.51E-12 |
| BF1210 | BF638R_1276 | *ahpC* | Alkyl hydroperoxide reductase subunit (O) | 77 | 76 | 93 | -1.0 | 9.86E-01 |
| BF2360 | BF638R_2372 | *tpx* | Thioredoxin peroxidase (O) | 7 | 12 | 10 | 1.8 | 8.22E-07 |
| BF2556 | BF638R_2545 | *sod* | Superoxide dismutase (P) | – | 8 | 4 | -1.2 | 4.54E-01 |
| BF2777 | BF638R_2786 | *tps* | Thiol peroxidase-scavengase (O) | – | 6 | – | -1.5 | 2.38E-02 |
| BF3656 | BF638R_3727 | *glp* | Glutathione peroxidase (O) | – | 14 | – | -1.4 | 3.75E-02 |
| BF2268 | BF638R_2327 | *dyp* | Dyp-type peroxidase (P) | – | 5 | 5 | -1.7 | 1.78E-04 |
| BF2362 | BF638R_2374 | *ccp1* | Cytochrome-C peroxidase (P) | 3 | – | – | 3.3 | 7.06E-28 |
| Metal metabolism |  |  |  |  |  |  |  |  |
| BF1317 | BF638R_1333 | *dps* | DNA and iron binding (P) | 25 | 38 | 45 | -3.4 | 6.02E-17 |
| BF2884 | BF638R_2891 | *ftnA* | FerritinA, non-haem iron (P) | – | 4 | – | 1.0 | 9.66E-01 |
| BF3271 | BF638R_3305 | *bfr* | Bacterioferritin (R) | – | 22 | – | -5.5 | 4.08E-25 |
| BF3032 | BF638R_3039 | *pir2* | Pirin (R) iron, cupin family | – | 15 | 6 | -2.3 | 2.13E-15 |
| BF3190 | BF638R_3217 | *nlpE* | Copper resistance; signalling for Cpx pathway (P) | – | 9 | – | 1.7 | 5.47E-06 |
| Nucleic acid repair |  |  |  |  |  |  |  |  |
| BF1191 | BF638R_1258 | *nfo* | Endonuclease IV (L) | – | 4 | – | -1.1 | 3.12E-01 |
| BF1780 | BF638R_4381 | *radC* | RecG-like activity (L) | – | – | 3 | -2.2 | 4.20E-04 |
| BF1945 | BF638R_1917 | *mgs1* | AAA-family ATPase, Holliday junction repair (L) | – | 4 | – | -1.1 | 4.16E-01 |
| BF2642 | BF638R_2636 | *uvrB* | UvrB SOS-repair system (L) | – | – | 3 | 1.2 | 2.30E-02 |
| BF3312 | BF638R_3340 | *ung* | Uracil-DNA glycosylase (L) | – | 6 | – | -2.7 | 6.00E-19 |
| BF4111 | BF638R_4168 | *ada* | Methylated-DNA –protein-cysteine methyltrans ( | – | 3 | – | 1.3 | 6.71E-03 |
| Protein repair/chaperones |  |  |  |  |  |  |  |  |
| BF1172 | BF638R_1237 | *clpA* | Chaperone ATPase (O) | – | 12 | – | -2.2 | 6.11E-08 |
| BF3207 | BF638R_3234 | *dnaJ* | DnaJ; chaperon for protein aggregates (O) | – | 10 | – | -2.2 | 1.07E-12 |
| BF3378 | BF638R_3405 | *msrA* | Methionine sulfoxide reductase (O) | – | 12 | 11 | -1.3 | 6.08E-03 |
| Redox balance |  |  |  |  |  |  |  |  |
| BF1279 | BF638R_1294 | *trxG* | Thioredoxin (O) | – | 14 | – | -2.8 | 5.54E-09 |
| BF2237 | BF638R_2295 | *trxD* | Thioredoxin (O) | 3 | 8 | 13 | 1.1 | 3.69E-01 |
| BF2656 | none | *trxX* | Thioredoxin-like no conserved cysteines (O) | – | 5 | – |  |  |
| BF2694 | BF638R_2701 | *trxC* | Thioredoxin-exported (O) | – | 7 | – | 1.2 | 4.47E-01 |
| BF3015 | BF638R_3022 | *trxE* | Thioredoxin (O) | – | 7 | 6 | -1.7 | 1.23E-04 |
| General function |  |  |  |  |  |  |  |  |
| BF2495 | BF638R_2455 | *uspA* | Universal stress protein (T) | – | 16 | 5 | -1.8 | 2.77E-04 |
| BF2778 | BF638R_2787 | *oim* | Membrane protein; HdeD domain, acid defence (S | – | 9 | – | -2.7 | 4.51E-08 |

<sup>a</sup> *B. fragilis* NCTC 9343 stress response genes upregulated during exposure to H<sub>2</sub>O<sub>2</sub>, air, or 5% oxygen (from Table 1 of (23)).
<sup>b</sup> Original *B. fragilis* NCTC 9343 locus tags.
<sup>c</sup> Cross-genome gene equivalency was determined by reciprocal best hit analysis using BLASTp. Locus tag names for *B. fragilis* 638R from RefSeq accession no. NC\_016776.1.
<sup>d</sup> COG classes: L, replication, recombination and repair; O, protein fate and chaperones; P, inorganic ion metabolism and transport; R, general function; S, no functional prediction; and T, signal transduction as reported (54).
<sup>e</sup> Fold induction as determined by ANOVA analysis of the Nimblegen microarray expression data. Only genes considered to be overexpressed relative to anaerobic conditions are enumerated, see ref (23).
<sup>f</sup> Fold change (FC) and adjusted *p*-values (*p* adj) as calculated by DESeq2 (51). Highlighted entries indicate significant (*p* < 0.001) differential expression of greater than 1.5-fold differential expression; red/green highlighting denotes over or under expression compared to anaerobic conditions, respectively.

Of the 550 previously identified essential genes of *B. fragilis* 638R (24) (494 of which were retained in the RefSeq genome annotation NC_016776.1) we identified 372 as core genes of *B. fragilis* (Table S1). Here we show that of those core essential genes, only 12 were significantly differentially regulated between the two conditions (Fig 1B). Three are more highly expressed under nanaerobic conditions, and nine are more highly expressed under anaerobic conditions (Fig 1B).

Next, we sought to determine whether a cellular process that requires large amounts of ATP is as active in bacteria grown nanaerobically. The Type VI secretion system is a dynamic inter-cellular antagonistic system that requires the assembly and disassembly of a toxin-loaded tube enclosed within a contractile sheath. After contraction of the sheath with subsequent propulsion of the tube across both membranes, the sheath is disassembled by ClpV (TssH). ClpV is an AAA+ ATPase (25) that uses the energy from ATP hydrolysis to mechanically disassemble the sheath structure (26). We found that *B. fragilis* 638R co-cultured under nanaerobic conditions with sensitive strain *B. vulgatus* CL09T02C10 shows similar T6SS killing activity as when cultured under anaerobic conditions (two log reduction of target strain under both conditions, Fig 1C). The RNASeq data revealed that nearly all genes of this T6SS locus are upregulated under nanaerobic conditions; however, only two of these genes reached significance by the criteria used in this study (Table S1). These data suggest that the GA3 T6SS of *B. fragilis* is functionally relevant at the oxygenated mucus layer as previously predicted (27, 28). These data collectively demonstrate that although there are transcriptional differences between anaerobic and nanaerobic growth, nanaerobically grown bacteria have similar transcriptional profiles of the vast majority of essential core genes as their anaerobically grown counterparts, are not undergoing general stress responses, and are metabolically and energetically active.

### Prevalence of cytochrome *bd* genes in the Bacteroidales

Bacteroidales is an order of bacteria that contains not only gut species of the genera *Bacteroides, Parabacteroides, Prevotella*, and *Alistipes*, but also oral pathogens such as *Porphyromonas gingivalis, Tannerella forsythia*, oral *Prevotella* species, as well as vaginal species and species that occupy other mammalian ecosystems. CydAB encoding genes were shown to be present in numerous *Bacteroides* species as well as *P. gingivalis* and *Prevotella intermedia* (2). In addition, a 2013 study using tBLASTx identified *cydAB* in 51 Bacteroidetes genomes (29); however, the species identifications within this large phylum were not provided. To determine the potential for nanaerobic growth of other species of Bacteroidales, nearly all of which associate with mammalian hosts, we queried a representative strain of 173 different species, contained within 32 genera and 13 families of Bacteroidales for pfams PF01654.17 and PF02322.15 specific to CydA and CydB, respectively. Protein sequences matching these models were detected in the genomes of 163 species (Table S2). Of the remaining 10 genomes, five contained *cydB*, three of which also had a *cydA* gene containing one or more frameshifts, possibly due to sequencing errors. These data strongly suggest that species of the order Bacteroidales that occupy diverse host ecosystems have the ability to respire aerobically under low oxygen conditions, transferring electrons from reduced quinone to oxygen *via* cytochrome *bd*, and will likely grow under nanaerobic conditions.

### Analysis of fluorescence of green and red fluorescent proteins from nanaerobically grown *Bacteroides* species

We sought to take advantage of this normal physiological property of these bacteria to determine if 0.10-0.14% oxygen is sufficient to mature GFP for imaging analyses. We used a plasmid created in the Sonnenburg lab, pWW3452, which contains an optimized promoter sequence and ribosome binding site resulting in very high GFP levels in *Bacteroides* (5) using the superfolder GFP gene (abbreviated as GFP throughout the paper) (30). This construct was created in the pNBU2-erm backbone vector that integrates into *Bacteroides* chromosomes (31). *Bacteroides* species containing pWW3452 were shown to fluoresce when grown anaerobically with subsequent exposure to oxygen for 60 minutes (5). We transferred pWW3452 into the chromosomes of four *Bacteroides* species and grew them under strict anaerobic conditions or in an atmosphere of 0.10 – 0.14% oxygen (1000 – 1400 ppm). After four hours of growth with shaking, cultures were added to agarose pads on microscope slides and sealed in their respective atmospheres in the chambers. Microscopic analysis revealed bright green fluorescence from bacteria grown nanaerobically but not anaerobically (Fig 2A, B). A similar analysis using confocal microscopy of the mCherry equivalent plasmid, pWW3515, revealed weak mCherry fluorescence when *B. fragilis* 638R was grown nanaerobically (Fig 2C). We attempted to improve the red fluorescence by analyzing three additional red fluorescent proteins, DsRed2, mKate2, and TagRFP. We replaced the GFP gene of pWW3452 with codon-optimized genes encoding each of these proteins (Fig S3) and transferred the resulting plasmids into *B. fragilis*. Histogram analysis of the images revealed that mKate2 produced the brightest fluorescence of the four red proteins (Fig 2C, D). Therefore, the chromophores of GFP, mCherry, and mKate2 mature to a significant level under nanaerobic conditions so that *Bacteroides* harboring them can be fluorescently imaged.

**Figure 2.**
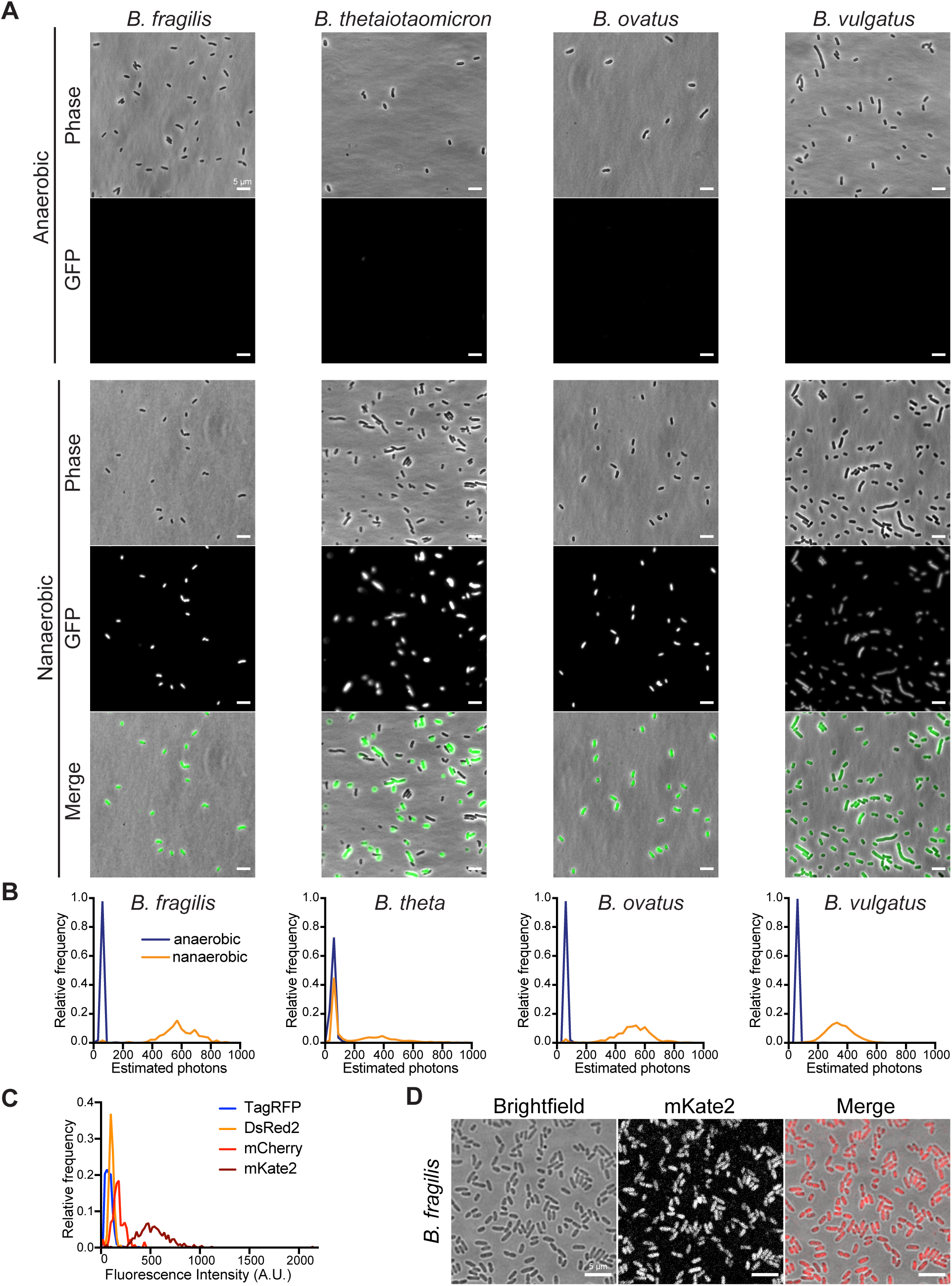
Nanaerobic growth allows fluorescent imaging of *Bacteroides* species. **A**. Epifluorescent microscopy of indicated *Bacteroides* species expressing GFP, grown under aerobic and nanaerobic conditions. Scale bars = 5 µm. **B**. Histograms of single-cell mean green signal intensity in cells of the indicated species expressing GFP. **C**. Histograms of red confocal fluorescence signal intensity in *B. fragilis* cells expressing the indicated red fluorescent protein grown under nanaerobic conditions. **D**. Confocal fluorescent microscopy of *Bacteroides fragilis* expressing mKate2, grown under nanaerobic conditions. Scale bars = 5 µm.

### GFP-labeling of proteins to track subcellular localization

To test whether we could use this process to label and track proteins in actively growing *Bacteroides*, we linked GFP to genes encoding two proteins secreted in outer membrane vesicles (OMVs). BACOVA_04502 of *B. ovatus* ATCC 8483 encodes an inulin lyase (32-34) and was previously shown to be secreted in OMVs by concentration of OMVs by two rounds of ultracentrifugation of culture supernatants followed by activity assays (33). To determine if we could visualize this protein in OMVs, we created a linker-GFP construct by altering pWW3452 (Fig 3A, C, D), fused the inulinase gene in frame, and transferred it to *B. ovatus*. Fluorescence microscopic analysis showed that the protein was present in the membrane and showed bright fluorescence in OMVs. Moreover, we could visualize inulinase-loaded OMVs vesiculating from the cell surface (Fig 3D). This is in stark contrast to the cytoplasmic localization of unlinked GFP shown in Fig 2B, and the lack of labeling of OMVs in these strains.

**Figure 3.**
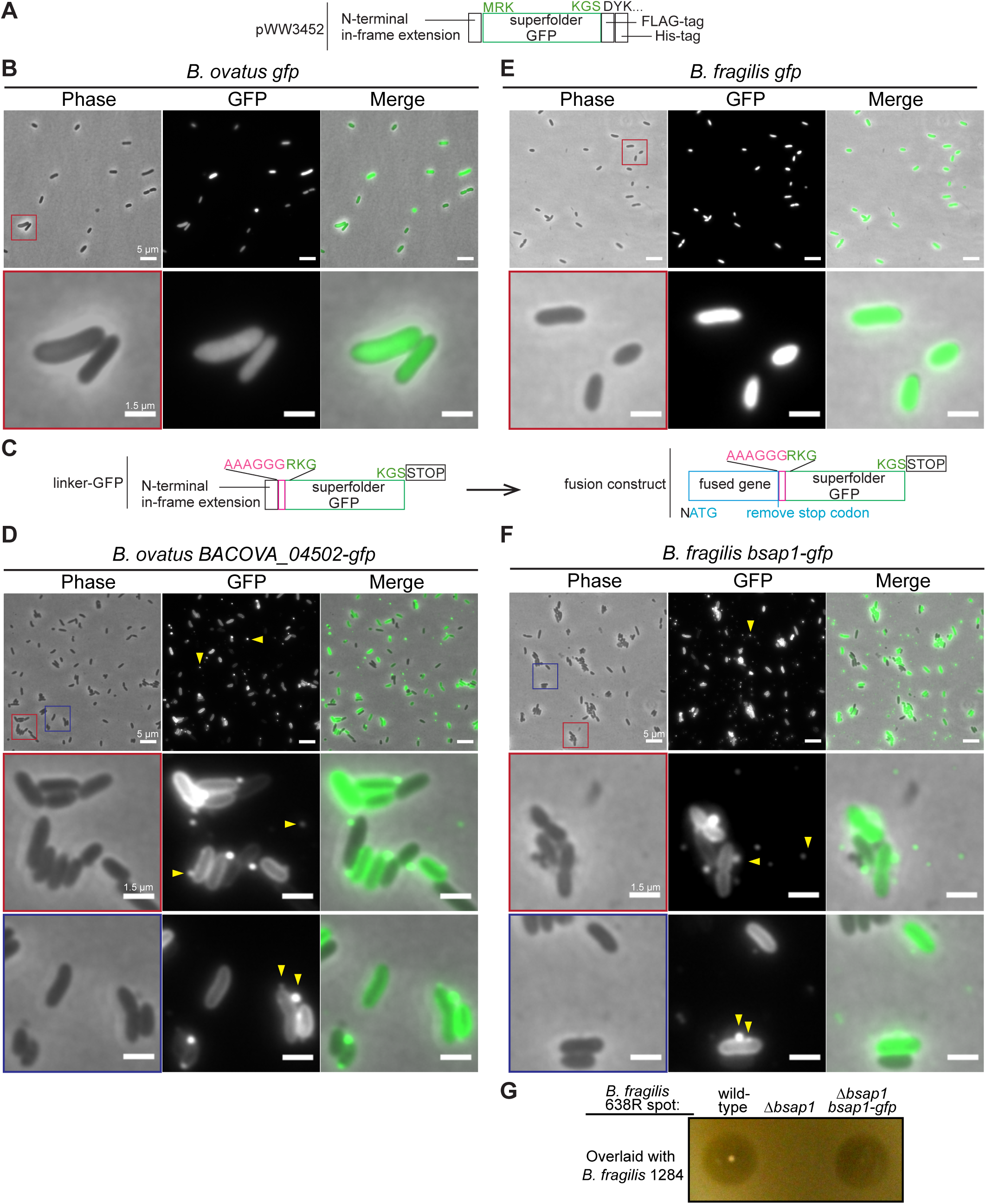
Localizaton of GFP-linked proteins in *B. fragilis*. **A**. Diagram of the superfolder GFP expression cassette of plasmid pWW3452 (5). **B, E**. *Bacteroides ovatus* and *B. fragilis* with pWW3452 display cytosolic localization of GFP. Second row corresponds to magnified field (scale bars = 1.5 µm) of the area indicated in red in the top row (scale bars = 5 µm). **C**. Modifications made to pWW3452 to create pLinkerGFP (left construct) and the addition of upstream genes to create fusion proteins with linker-GFP (right panel). N represents an additional nucleotide introduced to shift reading frame relative to N-terminal in-frame extension). **D**. *B. ovatus* expressing GFP-fused *BACOVA_04502*, encoding an inulinase, which is secreted in outer membrane vesicles (OMVs indicated with yellow arrows). Second and third rows correspond to magnified images (scale bars = 1.5 µm) of the areas indicated in red and blue in the top row (scale bars = 5 µm). **F**. *B. fragilis* expressing GFP-fused BSAP-1, which is abundantly secreted in OMVs (indicated with yellow arrows). Second and third rows correspond to magnified images (scale bars = 1.5 µm) of the areas indicated in red and blue in the top row (scale bars = 5 µm). **G**. Agar overlay assay of BSAP-1 sensitive strain *B. fragilis* 1284 to growth inhibition by the indicated *B. fragilis* 638R strains.

Similarly, we cloned into pLinkerGFP the BSAP-1 gene (BF638R_1646) of *B. fragilis*, encoding a MACPF domain antimicrobial toxin (35). BSAP-1 was previously shown to be secreted in OMVs by ultracentrifuging OMVs from culture supernatant and then performing immunofluorescence labeling using a primary antibody specific to BSAP-1 and a FITC-labeled secondary antibody (35). Here, we made the *bsap1-linker-gfp* fusion plasmid and integrated it into the chromosome of the BSAP-1 mutant strain. We first confirmed that the toxin was active with the C-terminal GFP fusion. The agar overlay shows that BSAP-1-GFP is able to restore antibacterial toxin activity to the *bsap-1* deletion mutant (Fig 3G). Using fluorescence microscopic analysis of the BSAP1-GFP fusion, we observed it in the membrane of the bacteria and also in OMVs (Fig 3F). As with the *B. ovatus* inulinase, instances of BSAP-1 in OMVs vesiculating from the cell were frequently observed. Similarly, fluorescence is absent in OMVs and membranes of the *B. fragilis* strain expressing the unlinked cytosolic GFP (Fig 3E). In sum, this process allows visualization and localization of two cytosolic proteins (unlinked GFP and mKate) and two OMV-associated proteins in live bacteria without the need to generate a specific antibody, perform an activity assay, or poison the cells by atmospheric oxygen exposure.

### Time-lapse microscopy reveals dynamic type VI secretion system processes in *B. fragilis*

Time-lapse microscopy of fluorescently-labeled T6SS structural components has played a key role in understanding the dynamics, regulation and mechanics of T6SS firing in many Proteobacterial species (reviewed (36)). *Bacteroides* species harbor three different architectures of T6SS, termed GA1-3 (37), which are largely still unexplored. *B. fragilis* 638R harbors a GA3 T6SS which has been shown to kill *Bacteroides* species under anaerobic (27, 38, 39) and in this study, under nanaerobic conditions (Fig 1C). We made a deletion mutant of BF638R_1993, encoding the GA3 T6SS protein TssB, one of two sheath proteins, which has been successfully fused to GFP to visualize sheath extension and contraction in Proteobacterial species growing in air (40). We cloned the *B. fragilis* 638R *tssB* into pLinkerGFP to synthesize a TssB-GFP fusion and introduced it into *B. fragilis* 638R Δ*tssB*. To confirm that the fusion to GFP did not abrogate TssB function, we tested whether the T6SS was still able to function using two different assays. First, we assayed for the presence of the tube protein, TssD, in the culture supernatant. There is no TssD tube protein in the supernatant of the Δ*tssB* mutant, and addition of the p*tssB-gfp* construct to the Δ*tssB* mutant restored TssD secretion (Fig 4A). In the second assay, we used antagonism of sensitive strain *B. vulgatus* CL09T03C10 as a readout for T6SS function. The complemented mutant restored the 2-log killing to the Δ*tssB* mutant, comparable to the level of antagonism by wild-type (Fig 4B). Therefore, as was demonstrated with the GFP fusion to the TssB protein in Proteobacterial species, this fusion does not interfere with the function of TssB.

**Figure 4.**
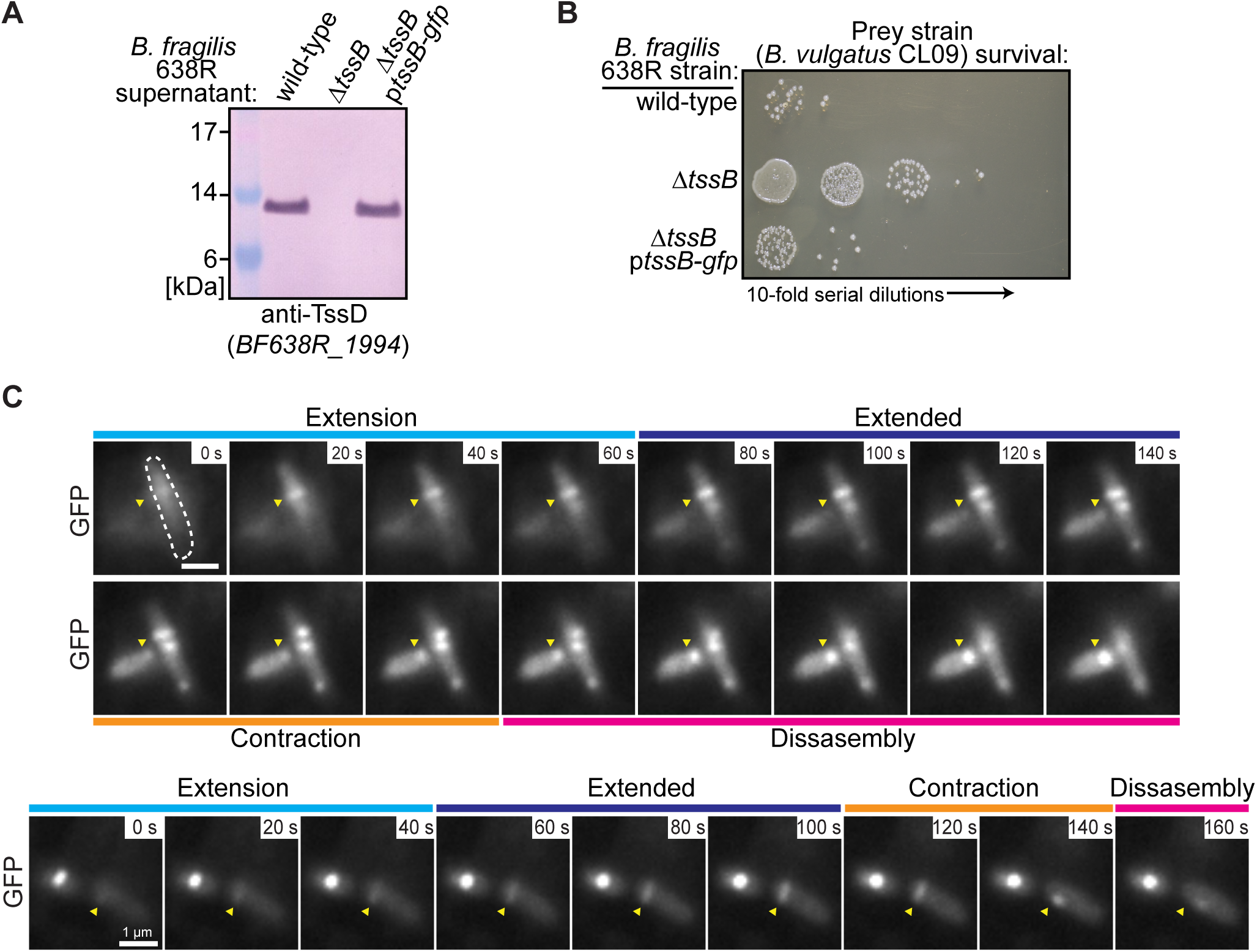
Fluorescence imaging of T6SS sheath dynamics in *B. fragilis*. **A**. Western immunoblot of culture supernatants from the indicated *B. fragilis* strains, probed with antiserum to TssD. **B**. T6SS antagonism assays with *B. fragilis* strains against the sensitive strain *B. vulgatus* CL09T03C10. Strains were co-cultured overnight under anaerobic or nanaerobic conditions and survival of the sensitive strain was quantified by serial 10-fold dilutions on selective media. **C**. Time-lapse of two cells undergoing T6SS firing. Yellow arrows indicate firing T6SS assemblies. Scale bars = 1 µm.

Next, we analyzed whether we could use the Δ*tssB*::p*tssB-gfp* strain to do live-cell time-lapse imaging of T6SS sheath assembly and contraction. Bacteria were grown nanaerobically for 4 hours, spotted on an agarose pad and sealed nanaerobically in a glass-bottom dish. Time-lapse imaging was conducted after a 15-20 minute incubation at 37ºC. TssB displayed diffuse cytosolic fluorescence in a large proportion of the population, and in many cells, nearly all the fluorescence was present in foci, indicative of a contracted sheath or sheath aggregates (Fig S4). Some cells had one or more extended sheaths (Fig 4 C, Fig S4, Supplemental video 1), usually along an axis perpendicular to the length of the cell and roughly as long as the cell width (∼ 0.5 µm). Sheath contraction was very rapid, going from a fully extended to fully contracted sheath between single frames (Fig S4 right panel, Fig 4C, Supplemental video 1). Upon contraction, sheath length changed to 0.2-0.3 um.

Visualization of the full sheath extension-contraction-disassembly cycle (Fig 4C, Supplemental video 1) indicates that multiple sheaths at different stages of assembly can occur in the same cell. Under the conditions in which these bacteria were imaged, sheath assembly can take up to 60 seconds, and the sheath can remain extended for more than four minutes before contracting (Supplemental video 1). Disassembly of contracted sheaths was highly variable between cells and lasted longer than one minute.

### Analysis of presence of cytochrome *bd* in other sequenced human gut bacteria

To determine if other gut bacteria currently known to grow only under anaerobic conditions may also be able to grow under nanaerobic conditions and therefore,may be amenable to fluorescent protein imagining analyses, we searched for protein sequences matching the same cytochrome *bd* (CydAB) profile HMM models used to query the Bacteroidales genome set. For this analysis, 612 genomes of the Human Gastrointestinal Bacteria Culture Collection (41) that contained protein translations and that were retrievable as assembled genomes from the NCBI databases were queried (Table S3). As shown above, the gut Bacteroidales uniformly contain cytochrome *bd*. There was variability in the phylum Actinobacteria, with some genera containing CydAB and others, such as *Bifidobacterium*, lacking them. *Eggerthella lenta* and several other species in the Eggerthellaceae family contain CydAB. Members of this species have been shown to inactivate the cardiac drug digioxin (42). Among the Firmicutes, many members of the class Bacilli are known to grow in the presence of oxygen, including many *Lactobacilli*, and most species in this class have CydAB. However, those in the class Clostridiales are strict anaerobes and lack CydAB. Among less studied Firmicutes, several genera in the class Negativicutes contain CydAB, including *Veillonella, Megamonas* and *Mitsuokella*. Therefore, in addition to the Bacteroidales and *Akkermansia*, there are potentially other gut bacteria currently classified as strict anaerobes that may be amenable to nanaerobic growth and the use of fluorescent protein analysis if gene transfer systems are available.

## DISCUSSION

The most important aspect of this study to the gut microbiome field is the application of nanaerobic growth to fluorescently image proteins and dynamic cellular processes in actively growing “nanaerobic” gut symbionts. In this regard, it was important to establish that growth under nanaerobic conditions is a normal process of these bacteria. Since the identification of nanaerobic respiration in *Bacteroides* in 2004 (2), there have been no further analyses of this process, and therefore, our understanding was very limited. Transcriptomic analysis of these two conditions has been very illuminating. Very few core essential genes are differentially expressed under the two conditions and there is no evidence of a general stress response during nanaerobic growth. In fact, many of the genes previously identified as upregulated during stress response to H_2_O_2_ or high levels of oxygen are more highly expressed under anaerobic conditions. A transcriptomic analysis of the anaerobic sulfate-reducing bacterium *Desulfovibrio vulgaris* showed a similar lack of upregulation of most of the oxidative stress response genes during growth at 0.1% oxygen compared to exposure to air (43). Oxygen is continually released from intestinal epithelial cells, creating an oxygen gradient that rapidly decreases from the mucus layer to the anaerobic lumen (1). As *Bacteroides* are present at both the mucus layer and in the lumen, it is not surprising that these bacteria have different transcriptional programs based on whether they find themselves in the presence or absence of nanaerobic concentrations of oxygen. It will be interesting to determine the functions of various gene products that are differentially regulated under each condition including the various transcriptional regulators, SusCD outer membrane nutrient transporters, and fimbriae genes, 11 of which are more highly expressed under anaerobic conditions.

There are numerous applications for the use of nanaerobic growth to fluorescently image these bacteria and their proteins, allowing for new types of analyses not easily performed using other methodologies. Such studies include the tracking of protein subcellular localizations and secretion. Here we show that we can readily localize proteins to OMVs, which have been shown in numerous studies to be loaded with molecules essential for interactions with other bacteria and the host. (33, 35, 44-48). Furthermore, as the chromophores of both red and green proteins mature under nanaerobic conditions, protein-protein interactions and co-localization studies in metabolically active bacteria are possible. Additionally, numerous types of single-cell analyses are possible to study transcriptional and phenotypic heterogeneity among a population of bacteria. Also, this process should allow for imaging of biofilm and multi-species structures in flow cells. Finally, the real-time imaging of *B. fragilis* GA3 T6SS sheath assembly and contraction demonstrates the ability of image dynamic cellular processes in these gut bacteria. In summary, for nanaerobic species such as *Bacteroides*, fluorescent protein imaging is now an available tool to explore numerous aspects of bacterial cell biology.

## METHODS

### Growth of bacterial strains

*Bacteroides* wild-type strains used in the study include *B. fragilis* 638R, *B. thetaiotaomicron* VPI-5482, *B. ovatus* ATCC 8483 and *B. vulgatus* ATCC 8482. *Bacteroides* strains were grown in BHI broth supplemented with 5g/L yeast extract and 5 µg/mL hemin (BHIS broth). Where appropriate, antibiotics were added: carbenicillin, 100 µg/mL; erythromycin, 5 µg/mL; gentamycin, 200 µg/mL; tetracycline, 6 µg/mL. *Bacteroides* were grown on BHIS plates which is BHI supplemented with 5 µg/mL hemin and 2.5 µg/mL vitamin K1. In some cases, modified M9 medium was used which is M9 minimal medium with 0.5% (wt/vol) glucose, cysteine to 0.025%, 5 µg/mL hemin, 2.5 µg/mL vitamin K1, 2 µg/ml FeSO_4_ 7H_2_O, and 5 ng/mL vitamin B_12_. For nanaerobic growth, oxygen and hydrogen levels were continually monitored using a CAM-12 gas monitor (Coy Lab Products, Grass Lake, Michigan).

### Creation of fumarate reductase (*frd)* mutation and complementation

The *frd* genes (BF638R_4499-4501) were deleted from the *B. fragilis* 638R (TM4000) genome by the following procedure. Regions flanking the deletion were PCR amplified with Phusion polymerase (NEB) and cloned directionally into the BamHI site of pLGB36 (49) using NEBuilder (NEB) and transformed into *E. coli* S17 λ pir. The plasmid was then conjugally transferred from *E. coli* to *B. fragilis* and cointegrates were selected on BHIS erythromycin plates. The cointegrate was passaged for five hours in non-selective medium and plated on BHIS plates containing 50 ng/mL anhydrotetracycline (aTC) to select for those that lost the plasmid by double crossover recombination. Deletion mutants were identified by PCR. For complementation studies, the three *frd* genes were amplified with their native promoter using the primers listed in Table S4 and cloned into the BamHI site of pNBU2. This plasmid was introduced into Δ*frd* by conjugal transfer from *E. coli* S17 λ pir. All plasmids created for this study were sequenced at the Massachusetts General Hospital DNA Core Facility (Boston, MA).

### Growth analyses of *Bacteroides* species under anaerobic and nanaerobic conditions

All nanaerobic growth in this study was performed at 37ºC in an atmosphere of 1000-1400 ppm oxygen (0.10 – 0.14%) with a hydrogen concentration between 2.5 – 3.0%, with 10% CO_2_ and the remainder nitrogen. The same gas mix was used for the anaerobic conditions without oxygen. Cultures were grown to late log phase in BHIS under either anaerobic or nanaerobic conditions in shaking 5 mL cultures in 100 mL flasks. Aliquots of 3 µL were added to triplicate wells containing 100 µL of BHIS in the outer wells of a 96-well plate and growth was recorded over time at OD_600_ using an Eon high-performance microplate spectrophotometer (BioTek Instruments) with constant shaking between readings. Each of the growth curves summarizes 12 hours of growth, with OD_600_ readings taken every five minutes. Each curve of Fig 1 plots at one-hour intervals the average reading from between two and six replicates of each strain, with a Gompertz growth least squares fit line connecting them. The logarithmic-scaled graphs in Fig S1 are an alternate representation of the same data, except the smoothed lines were generated by locally weighted scatterplot smoothing (lowess) regression analysis.

### RNAseq analysis

Three biological replicate cultures of *Bacteroides fragilis* 638R were grown under anaerobic and nanaerobic conditions (5 mL) in 25 mL vented-top cell culture flasks with shaking for seven hours until an OD_600_ of 0.6 was reached. One mL of culture was added to microfuge tubes and sealed in their respective atmospheres and the bacteria were collected by centrifugation. The supernatant was quickly removed and the tube was immediately plunged into a dry ice ethanol bath. RNA extraction, library preparation, rRNA depletion, and sequencing (2 × 150 bp, Illumina HiSeq) were performed by Genewiz (South Plainfield, NJ). The sequencing reactions of the six samples each produced between 50.9 and 56.1 million reads (15.3 to 16.8 Mbases) with mean quality scores of 35.8 – 35.9 for each sample; all samples had ∼93% of bases with a quality score of greater than 30. The RNA-Seq data returned from Genewiz was evaluated using FastQC (v0.11.9, http://www.bioinformatics.babraham.ac.uk/projects/fastqc/) BBDuk (version 38.76, https://sourceforge.net/projects/bbmap/) was used to remove adapter sequences and to quality trim the raw Illumina reads. Reads with less than 50 bp remaining after QC and adapter trimming were discarded.

The cleaned reads were mapped to the *B. fragilis* 638R genome sequence contained in RefSeq accession no. NC_016776.1 using EDGE-pro (50), chosen because it was developed and optimized for mapping reads to prokaryotic sequences. Per-feature read mapping counts produced by EDGE-pro were evaluated for differential expression using both DESeq2 (51) and edgeR (52). For our analysis, a differentially expressed gene was defined as one that was determined to be expressed on average at least 1.5 fold up or down under nanaerobic conditions relative to anaerobic conditions by both DESeq2 and edgeR, and whose adjusted *p*-value (DESeq2) and FDR (edgeR) both were less than or equal to 0.001.

Among the essential genes of *B. fragilis* 638R previously reported (24), we identified “core” essential genes from this list as those whose products are similar at 90% or greater *via* blastp to gene products in five other *B. fragilis* strains analyzed: NCTC 9343, YCH46, CL03T12C07, CL05T12C13, and CL07T12C05. Volcano plots were created in Prism version 8.4.2 (GraphPad Software, San Diego, CA). BAM files containing the sorted mapping results of EDGE-pro have been deposited to the NCBI SRA database and assigned accession number PRJNA630209.

### Analysis for the presence of CydAB in the genomes of Bacteroidales and diverse human gut symbionts

Two sets of bacteria genomes were retrieved for these analyses. First, a representative set of 173 Bacteroidales genomes was downloaded from NCBI using Entrez queries (Table S2). In addition, a 612 genome subset of the Human Gastrointestinal Bacteria Culture Collection (41) that comprises assembled genomes with annotation and protein translations was retrieved from either the GenBank or RefSeq NCBI databases. Profile HMM models PF01654.17 (Cyt_bd_oxida_I) and PF02322.15 (Cyt_bd_oxida_II) were extracted from the Pfam 32.0 database (53) and compressed into binary form using the hmmpress program from the HMMER version 3.3 set of utilities (http://hmmer.org/). The proteomes were searched using hmmsearch utilizing the gathering threshold cut-off for each model.

### COG analysis

The *B. fragilis* 638R proteome contained in RefSeq accession NC_016776.1 was mapped to COG categories (54) using version 2.0.1 of the stand-alone eggNOG-mapper program (55) and version 5 of the EggNOG database (56) under default settings. Graphs demonstrating the RNA-Seq expression levels of these sets of genes were created in Graphpad Prism version 8.4.2, plotting the least significant fold-change/adjusted p-value for each as calculated by either DESeq2 or edgeR.

### Creation of red fluorescent protein plasmids

The genes for the red fluorescent proteins, DsRed2, mKate2, and TagRFP were codon optimized for *Bacteroides* (Fig S3) and synthesized by Genscript. Each gene was amplified with the primers listed in Table S4 and used to replace the super folder GFP gene and the downstream Flag- and His-tags of plasmid pWW3452 using NEBuilder (Fig 3D). Plasmids were transferred from *E. coli* S17 λ pir to *B. fragilis* 638R by conjugation.

### Creation of linker-*gfp* vector (pLinkerGFP) for generation of fusion proteins

A 753-bp piece of DNA was synthesized by GenScript containing the superfolder GFP open reading frame of pWW3452 (5) including a stop codon without the downstream Flag and His-tags and including the upstream liner region GCAGCTGCAGGAGGTGGA encoding Ala-Ala-Ala-Gly-Gly-Gly used by Basler *et al*. (40) for the fusion of proteins to *gfp*. This DNA was amplified with primers linker-GFP-F + R (Table S4). Primers pWW3452 linker GFP_F + R were used to amplify the remainder of pWW3452 lacking the GFP gene. These two parts were amplified using Phusion polymerase (NEB) and joined using NEBuilder (NEB). The plasmid (pLinkerGFP) was sequenced to validate its proper construction. As with the parent pWW3452, the β-lactamase gene is split, which does not affect its ability to confer resistance to ampicillin or carbenicillin. Genes cloned into pLinkerGFP for fusions with GFP at the C-terminus should include a single nucleotide upstream of the start codon of the cloned gene to make it out of frame with a portion of an open reading frame upstream in the regulatory region of pWW3452 (Fig 3D).

### Creation of fusion proteins with GFP in pLinkerGFP

Three different genes were cloned into pLinkerGFP to place the cloned genes upstream of the linker-GFP. BF638R_1646 (*basp-1*) was amplified from *B. fragilis* 638R, BACOVA_04502 was amplified from *B. ovatus* ATCC 8483, and BF638R_1993 (*tssB*) was amplified from *B. fragilis* 638R using primers listed in Table S4. All genes had an extra nucleotide added before the start codon to place the gene out of frame with a portion of an open reading frame upstream in the vector. For all clones, the pLinkerGFP vector was amplified with primers pLinkerGFP_F + R. Each gene was joined with the amplified pLinkerGFP using NEBuilder and transformed into *E. coli* S17 λ pir. All plasmids were subject to whole plasmid sequencing to confirm the correct joining of the segments. Clones were mated from *E. coli* into the appropriate *Bacteroides* strain with subsequent integration into tRNA^ser^ att site.

### BSAP-1 overlay assay

Bacterial cultures tested for BSAP-1 production were dotted onto plates (5 µl) and grown for 12 hours. The bacterial growth was removed with a swab and the plates were exposed to chloroform vapor for 15 minutes. BHIS top agar (4 ml) was inoculated with *B. fragilis* strain 1284 (BSAP-1 sensitive strain) and applied to the plate and zones of inhibition were imaged after nine hours.

### Western blot analyses

Antiserum to purified His-BF638R_1994 (TssD) was previously described (27). Supernatants from overnight bacterial cultures were boiled in LDS sample buffer and separated by electrophoresis using NuPAGE 12% polyacrylamide gels (Life Technologies). The contents of the gels were transferred to PVDF membranes, blocked with skim milk, and probed with (α-TssD) followed by alkaline phosphatase-labeled α-rabbit IgG, and developed with BCIP/NBT (KPL).

### T6SS antagonism assays

Strains included *B. fragilis* 638R, isogenic mutants ΔT6SS (27), Δ*tssB*, or Δ*tssB*p*tssB-gfp*. Log phase cultures of these strains were mixed at a ratio of 10:1 (v:v) with log-phase *B. vulgatus* CL09T04C04 as the sensitive strain. A total of 10 µL of the above mixtures were spotted on BHIS plates and incubated overnight in either an anaerobic or nanaerobic atmosphere. The spots were excised, suspended in PBS and serial 10-fold dilutions were plated to BHIS containing tetracycline (6 µg/mL) to select for *B. vulgatus* CL09T03C04.

### Widefield microscopy analysis of *Bacteroides* species grown anaerobically and nanaerobically

Bacteria were swabbed from a BHIS erythromycin plate directly into modified M9 medium pre-incubated overnight in anaerobic or nanaerobic atmosphere with erythromycin and grown shaking at 140 rpm for at least four hours. 10 µL of the bacteria were dotted onto 1.5% agarose pads made with modified M9 medium, a coverslip was added and sealed under nanaerobic or anaerobic conditions. Images were obtained using a Nikon Ti inverted microscope equipped with a Nikon motorized stage, an Andor Zyla 4.2 Plus sCMOS camera, Lumencore SpectraX LED Illumination, Plan Apo lambda 100×/1.45 NA Oil Ph3 DM objective lens, and Nikon Elements 4.30 acquisition software. The green channel was imaged using a Chroma 49002 filter cube. Images were adjusted and cropped using Fiji (57). Cell segmentation analyses and quantification of mean single-cell fluorescence signal intensity were carried out using Microbe J (58). The conversion factor from greys to estimated photons was computed using the formula (Full well capacity)/((max intensity)-offset) and corresponded to 0.28 estimated photons/intensity level, based on camera settings of 12 bit low gain (Full well capacity =1,100 e-; max intensity 4095 intensity levels; camera offset: ∼110 grey levels). Fluorescence intensity histograms were computed and plotted in Prism 8.

For time-lapse microscopy of T6SS firing, cells were grown as described above and 10 µl of each strain were spotted onto 1.5% agarose pads (800 µL, 22 × 22 mm) made with modified M9 medium. Pads were transferred to sterile glass-bottom dishes (Greiner Bio-One CELLview), tightly sealed with anaerobic vinyl tape (Coy Laboratory Products, Inc., Grass Lake, MI) and kept at 37ºC for 15-20 minutes. Cells were imaged as described above, with 100 ms exposure time in the green channel and 10 or 20-second intervals between images. The phase and red channels (Fig S4) were only imaged at the first time-point, using a Chroma 49008 filter cube for the red channel. Images were analyzed and the video was made using Fiji (57). Photobleaching in the green channel was corrected using the Bleach Correction tool with Histogram Matching. For the video, crops were upscaled using bilinear interpolation.

### Confocal microscopy of *B*. *fragilis* expressing different red fluorescent proteins

Cells were grown nanaerobically and spotted on sealed agarose pads as described above, and imaged on a Leica SP8 inverted confocal microscope equipped with a motorized stage, an HyD detector, lasers Kit WLL2 from 470nm to 670nm, HC PL APO 100×/1.45 oil objective lens, and LAS-X acquisition software. The imaging conditions were 1 Airy unit pinhole size, scan rate 400 Hz, line averaging 4. Excitation was carried out at 565 nm and emission collected at 610-670 nm. Laser settings were adjusted for maximum signal but were kept constant for all images. Transmitted light was collected using the transmitted standard PMT. Images were analyzed and quantified as described previously for GFP, without conversion from intensity levels to estimated photons.

## ACKNOWLEDGEMENTS

We thank Ryan Stephansky and Tim Ross-Elliott of the MicRoN Imaging facility at Harvard Medical School for technical assistance, and Sergey Pryshchep at the RPI Center for Biotechnology and Interdisciplinary Studies Microscopy Core (NSF grant # NSF-MRI-1725984). The Nikon Ti microscope at the MicRoN facility was purchase with the S10 shared equipment grant RR027344-0. MB is supported by SNSF Starting Grant BSSGI0_155778. This work was funded by Public Health Service grant R01AI132580 to LEC, MHM and BB and R01AI120633 to LEC from the NIH/NIAID. The funders had no role in study design, data collection and interpretation, or the decision to submit the work for publication.

## Author contributions

LGB designed research, performed research, analyzed data and wrote the paper. MJC analyzed data and wrote the paper, NH, SV and TI performed research, PML and MB designed research and analyzed data, MHM designed research, BB designed research, performed research, analyzed data, LEC designed research, performed research, analyzed data and wrote the paper.

---

**Figure S1**. Growth curves of Figure 1A plotted in log scale.

**Figure S2**. Volcano plots of RNAseq data separated into functional (*susCD*) or COG categories. Green dots represent genes down regulated under nanaerobic conditions and red dots represent genes upregulated under nanaerobic conditions. All data are contained in Table S1.

**Figure S3**. DNA sequences of the *Bacteroides* codon-optimized Ds-Red2, Tag-RFP and mKate2 genes used in Figure 2.

**Figure S4**. Epifluorescence microscopy image of *B. fragilis* 638R Δ*tssB* expressing a TssB-GFP fusion protein, spotted in a pad with *B. thetaiotaomicron* expressing mCherry. Scale bars = 5 µm in left panel and 1 µm in right panel. Yellow arrows in left panel indicate examples of extended sheaths. White arrow in the right panel indicates an extended T6SS sheath contacting to form a focus.

**Supplemental video 1**. Time-lapse of multiple cells of *B. fragilis* 638R Δ*tssD ptssB-gfp*. White lines indicate outlines of cells with a dynamic T6SS sheath.

