## Supplementary figures and images for "Nanaerobic growth enables direct visualization of dynamic cellular processes in human gut symbionts"

### supplemental figure 1

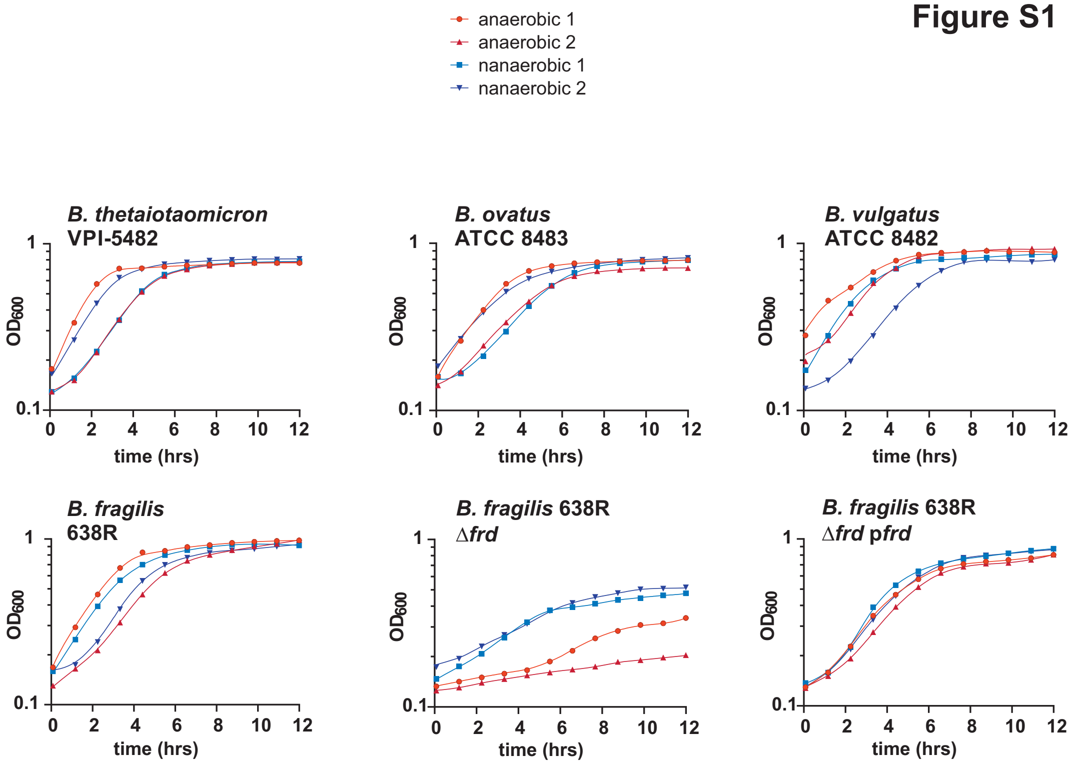

### supplemental figure 2

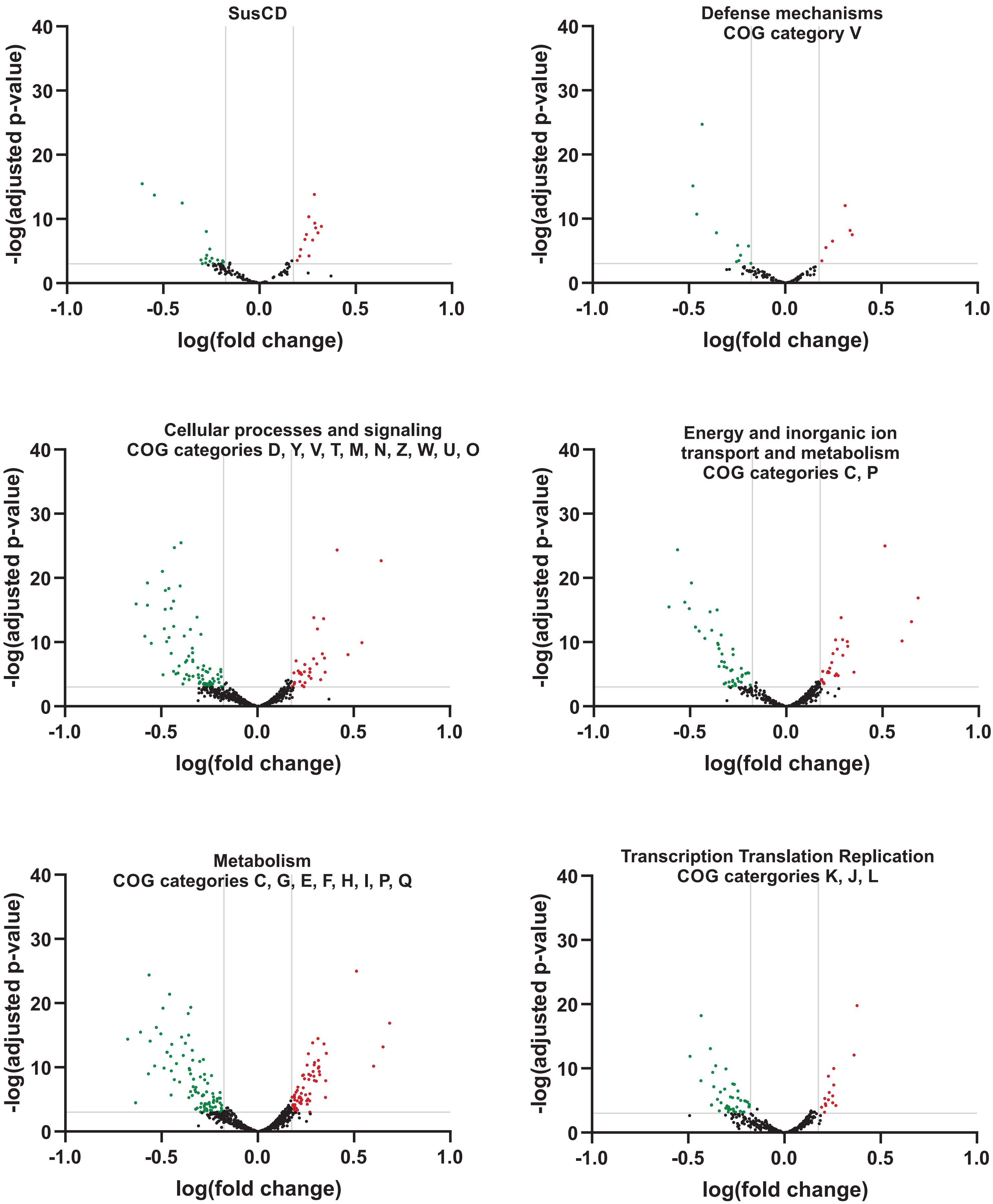

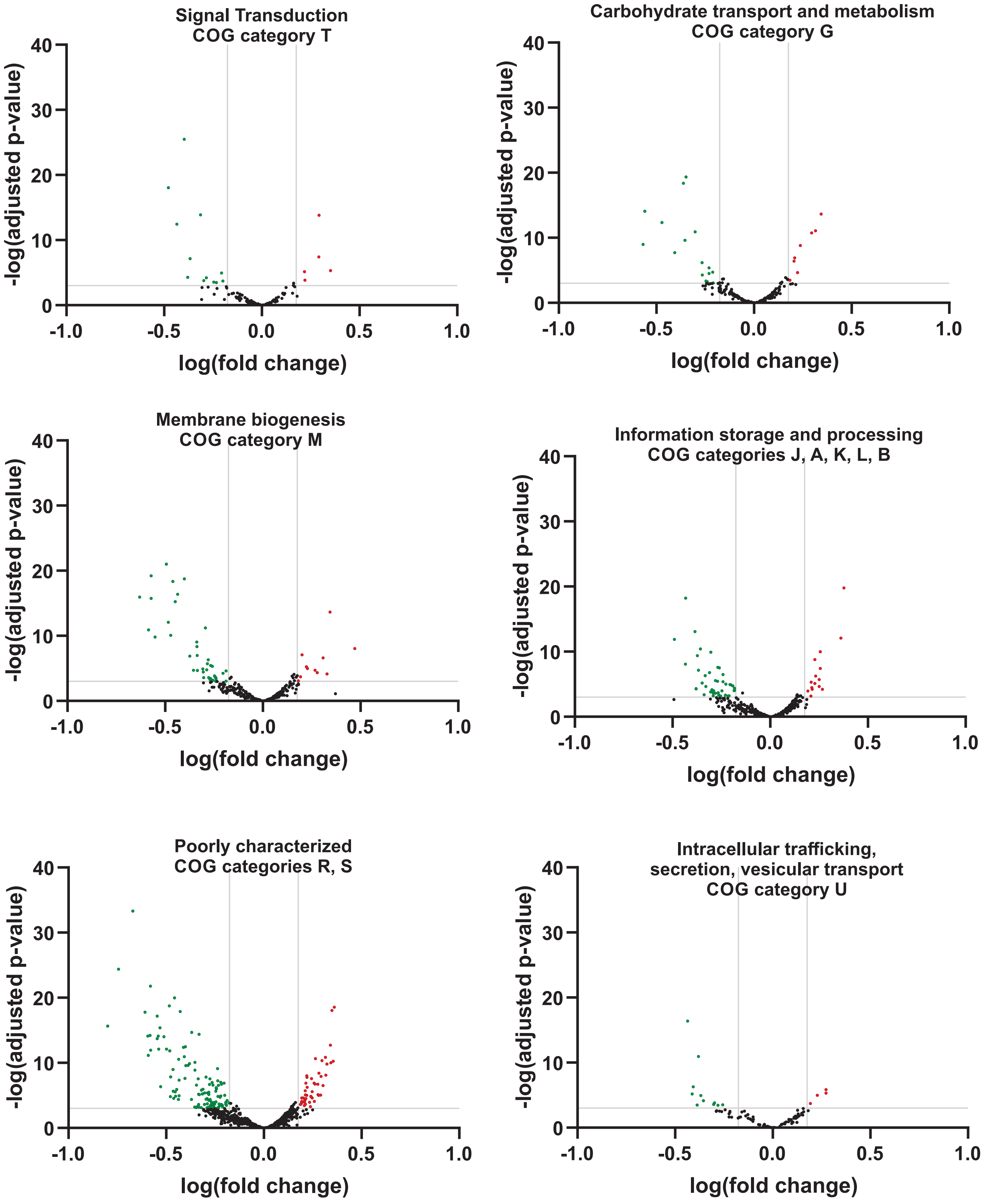

### supplemental figure 4

#### Figure S4

*B. fragilis* 638R ΔtssB-p*tssB*-gfp vs. *B. thetaiotaomicron* mCherry

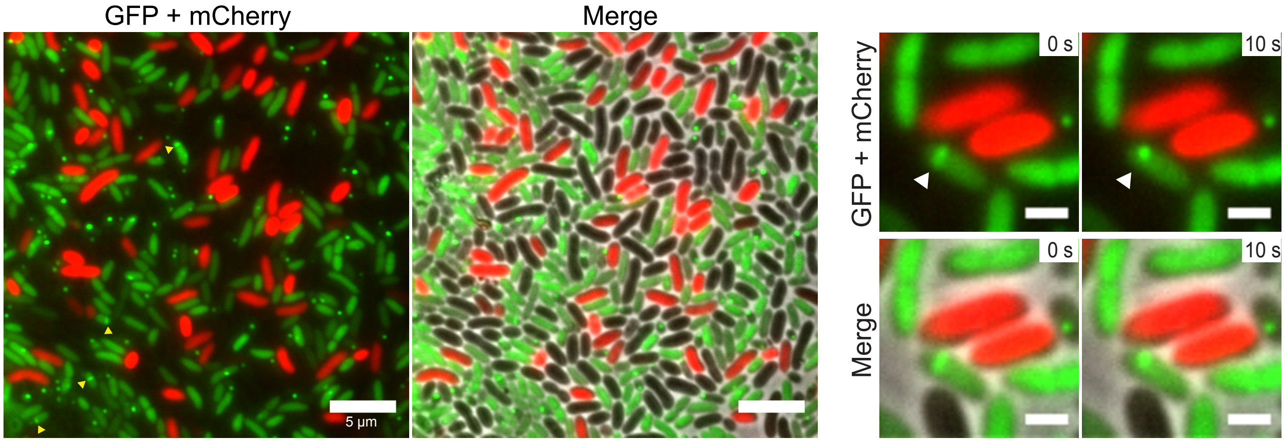
