## supplemental figure 3 for "Nanaerobic growth enables direct visualization of dynamic cellular processes in human gut symbionts"

### Figure S3. Sequence of fluorescent red protein genes codon optimized for *Bacteroides*

Gene name: DsRed2
Length:678bp
Sequence:
ATGGCCTCTAGTGAAAACGTAATCACAGAATTCATGCGTTTCAAA
GTACGTATGGAAGGTACCGTGAATGGACATGAATTTGAAATCGAA
GGAGAAGGTGAAGGACGTCCGTATGAAGGACATAACACAGTAAAG
CTGAAGGTGACCAAAGGTGGACCGCTGCCTTTTGCTTGGGATATT
TTGTCTCCGCAGTTCCAATATGGTAGTAAGGTATATGTGAAGCAT
CCGGCAGATATCCCTGATTATAAGAAGCTGTCTTTCCCTGAAGGA
TTCAAATGGGAACGTGTAATGAATTTTGAAGATGGAGGAGTAGCT
ACCGTGACTCAGGATTCGAGCTTGCAAGATGGTTGTTTCATCTAT
AAAGTAAAGTTCATCGGTGTGAATTTCCCGTCGGATGGACCTGTG
ATGCAGAAGAAAACCATGGGATGGGAAGCAAGCACAGAACGTCTG
TATCCTCGTGATGGTGTATTGAAAGGAGAAACTCATAAAGCCCTG
AAATTGAAAGATGGAGGACATTATTTGGTGGAATTCAAGTCTATC
TATATGGCCAAAAAACCGGTACAGCTGCCTGGTTATTATTATGTG
GATGCTAAGTTGGATATCACCAGTCATAACGAAGATTATACTATC
GTGGAACAATATGAACGTACCGAAGGACGTCATCATTTGTTTCTG
TAA

Gene name: mKate2
Length: 699bp
Sequence:
ATGGTATCTGAACTGATCAAGGAAAACATGCATATGAAGTTGTAT
ATGGAAGGTACCGTGAACAACCATCATTTCAAGTGTACTAGTGAA
GGTGAAGGAAAACCGTATGAAGGAACTCAGACAATGCGTATTAAA
GCTGTAGAAGGTGGACCGCTGCCTTTTGCTTTCGATATCTTGGCA
ACCTCTTTCATGTATGGTAGTAAGACTTTCATCAACCATACACAG
GGAATCCCGGATTTCTTTAAACAATCGTTTCCTGAAGGTTTCACA
TGGGAACGTGTAACCACTTATGAAGATGGAGGAGTACTGACCGCC
ACTCAGGATACCAGCCTGCAAGATGGATGTTTGATCTATAACGTA
AAGATCCGTGGTGTGAACTTCCCGTCGAATGGACCTGTAATGCAG
AAGAAAACCTTGGGTTGGGAAGCCAGCACAGAAACCCTGTATCCG
GCTGATGGAGGACTGGAAGGACGTGCAGATATGGCCCTGAAACTG
GTAGGTGGAGGACATCTGATCTGTAACTTGAAGACAACCTATCGT
TCTAAAAAACCGGCAAAGAACCTGAAGATGCCTGGTGTATATTAT
GTGGATCGTCGTTTGGAACGTATCAAAGAAGCCGATAAAGAAACA
TATGTGGAACAACATGAAGTAGCCGTGGCTCGTTATTGTGATCTG
CCTTCTAAATTGGGACATCGTTAA

Gene name: TagRFP
Length: 714bp
Sequence:

ATGGTATCGAAAGGTGAAGAATTGATCAAGGAAAACATGCATATG
AAACTGTATATGGAAGGAACCGTGAACAACCATCATTTCAAGTGT
ACTAGCGAAGGTGAAGGAAAACCGTATGAAGGAACTCAGACAATG
CGTATTAAAGTAGTGGAAGGTGGACCGTTGCCTTTTGCATTCGAT
ATCCTGGCCACTTCTTTCATGTATGGTAGTCGTACATTCATCAAT
CATACCCAGGGAATCCCGGATTTCTTTAAGCAATCTTTCCCTGAA
GGTTTCACATGGGAACGTGTAACCACTTATGAAGATGGAGGAGTA
TTGACCGCAACTCAGGATACCTCGTTGCAAGATGGATGTCTGATC
TATAACGTAAAGATCCGTGGTGTGAACTTTCCGAGCAATGGACCT
GTAATGCAGAAGAAAACCTTGGGTTGGGAAGCTAATACAGAAATG
TTGTATCCGGCAGATGGAGGACTGGAAGGACGTTCTGATATGGCC
CTGAAACTGGTAGGTGGAGGACATCTGATCTGTAACTTCAAGACA
ACCTATCGTAGTAAAAAACCGGCCAAGAACTTGAAGATGCCTGGT
GTATATTATGTGGATCATCGTCTGGAACGTATCAAAGAAGCTGAT
AAAGAAACCTATGTAGAACAACATGAAGTAGCCGTGGCTCGTTAT
TGTGATCTGCCTTCTAAATTGGGACATAAACTGAATTAA
